# Virological characteristics of the SARS-CoV-2 XEC variant

**DOI:** 10.1101/2024.10.16.618773

**Authors:** Yu Kaku, Kaho Okumura, Shusuke Kawakubo, Keiya Uriu, Luo Chen, Yusuke Kosugi, Yoshifumi Uwamino, MST Monira Begum, Sharee Leong, Terumasa Ikeda, Kenji Sadamasu, Hiroyuki Asakura, Mami Nagashima, Kazuhisa Yoshimura, The Genotype to Phenotype Japan (G2P-Japan) Consortium, Jumpei Ito, Kei Sato

## Abstract

The SARS-CoV-2 JN.1 variant (BA.2.86.1.1), arising from BA.2.86.1 with spike protein (S) substitution S:L455S, outcompeted the previously predominant XBB lineages by the beginning of 2024. Subsequently, JN.1 subvariants including KP.2 (JN.1.11.1.2) and KP.3 (JN.1.11.1.3), which acquired additional S substitutions (e.g., S:R346T, S:F456L, and S:Q493E), have emerged concurrently. As of October 2024, KP.3.1.1 (JN.1.11.1.3.1.1), which acquired S:31del, outcompeted other JN.1 subvariants including KP.2 and KP.3 and is the most predominant SARS-CoV-2 variant in the world. Thereafter, XEC, a recombinant lineage of KS.1.1 (JN.13.1.1.1) and KP.3.3 (JN.1.11.1.3.3), was first identified in Germany on August 7, 2024. XEC acquired two S substitutions, S:T22N and S:F59S, compared with KP.3 through recombination, with a breakpoint at genomic position 21,738–22,599. We estimated the relative effective reproduction number (Re) of XEC using a Bayesian multinomial logistic model based on genome surveillance data from the USA, the United Kingdom, France, Canada, and Germany, where this variant has spread as of August 2024. In the USA, the Re of XEC is 1.13-fold higher than that of KP.3.1.1. Additionally, the other countries under investigation herein showed higher Re for XEC. These results suggest that XEC has the potential to outcompete the other major lineage including KP.3.1.1. We then assessed the virological properties of XEC using pseudoviruses. Pseudovirus infection assay showed that the infectivity of KP.3.1.1 and XEC was significantly higher than that of KP.3. Although S:T22N did not affect the infectivity of the pseudovirus based on KP.3, S:F59S significantly increased it. Neutralization assay was performed using three types of human sera: convalescent sera after breakthrough infection (BTI) with XBB.1.5 or KP.3.3, and convalescent sera after JN.1 infection. In all serum groups, XEC as well as KP.3.1.1 showed immune resistance when compared to KP.3 with statistically significant differences. In the cases of XBB.1.5 BTI sera and JN.1 infection sera, the 50% neutralization titers (NT50s) of XEC and KP.3.1.1 were comparable. However, we revealed that the NT50 of XEC was significantly (1.3-fold) lower than that of KP.3.1.1. Moreover, both S:T22N and S:F59S significantly (1.5-fold and 1.6-fold) increased the resistance to KP.3.3 BTI sera. Here we showed that XEC exhibited higher pseudovirus infectivity and higher immune evasion than KP.3. Particularly, XEC exhibited more robust immune resistance to KP.3.3 BTI sera than KP.3.1.1. Our data suggest that the higher Re of XEC than KP.3.1.1 is attributed to this property and XEC will be a predominant SARS-CoV-2 variant in the world in the near future.

## Text

The SARS-CoV-2 JN.1 (BA.2.86.1.1) variant, arising from BA.2.86.1 with spike protein (S) substitution S:L455S, outcompeted the previously predominant XBB lineages by the beginning of 2024.^1^ Subsequently, JN.1 subvariants including KP.2 (JN.1.11.1.2) and KP.3 (JN.1.11.1.3), which acquired additional S substitutions (e.g., S:R346T, S:F456L, and S:Q493E), have emerged concurrently (**Figure 1A**).^2,3^ As of October 2024, KP.3.1.1 (JN.1.11.1.3.1.1), which acquired S:31del, outcompeted other JN.1 subvariants including KP.2 and KP.3, and is the most predominant SARS-CoV-2 variant in the world.^4^

**Figure 1.**
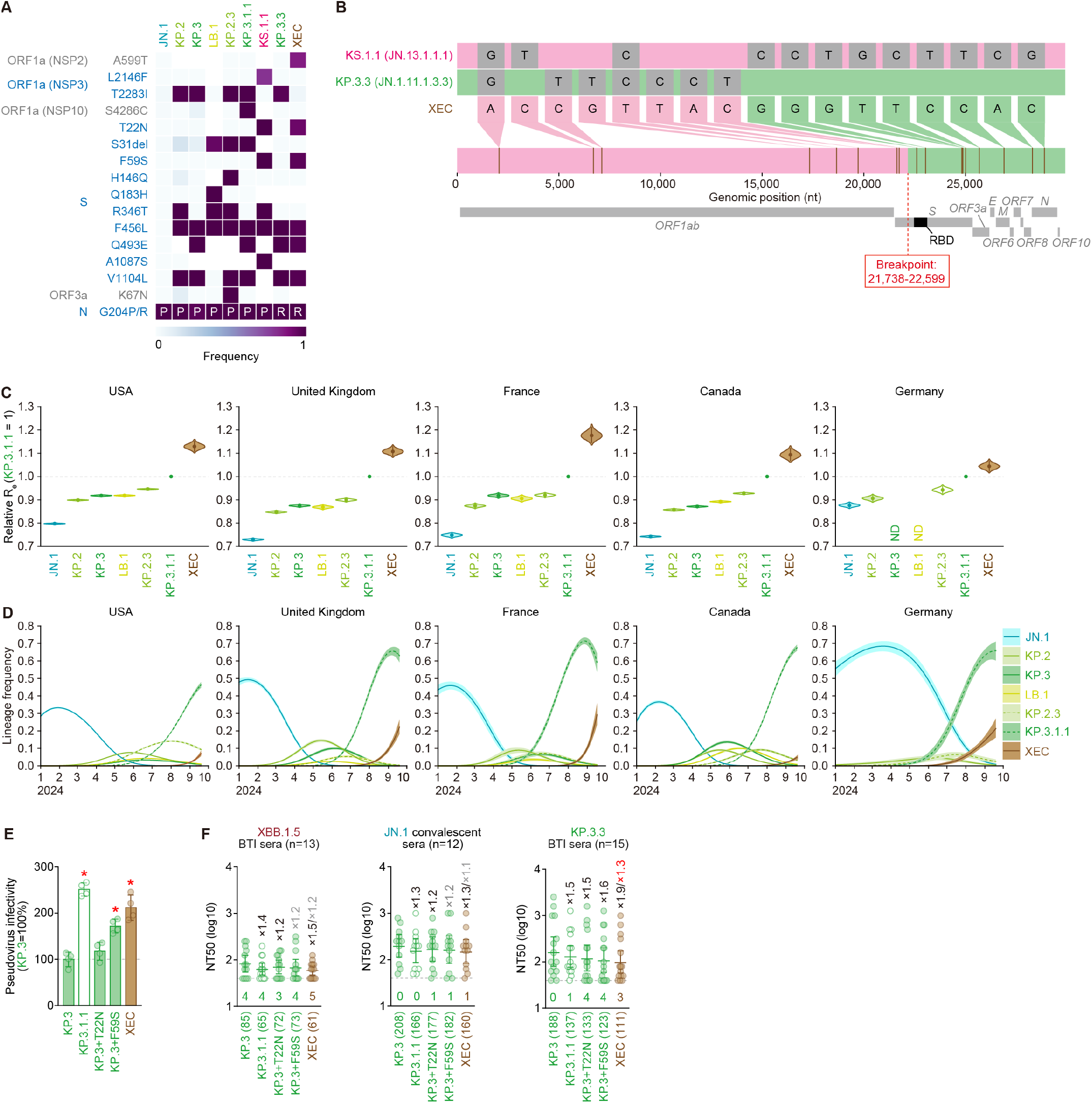
Virological features of XEC. **(A)** Frequency of mutations in XEC, and other lineages of interest. Only mutations with a frequency >0.5 in at least one but not all the representative lineages are shown. **(B)** Nucleotide differences between the consensus sequences of the KS.1.1, KP.3.3 lineages (parental lineages of XEC) and the XEC lineage, highlighting the recombination breakpoint. The plot was visualized with snipit (https://github.com/aineniamh/snipit). **(C)** Estimated relative R_e_ of the variants of interest in the USA, the United Kingdom, France, Canada, and Germany. The relative R_e_ of KP.3.1.1 is set to 1 (horizontal dashed line). Violin, posterior distribution; dot, posterior mean; line, 95% credible interval. **(D)** Estimated epidemic dynamics of the variants of interest in in the USA, the United Kingdom, France, Canada, and Germany from January 1, 2024, to September 19, 2024. Countries are ordered according to the number of detected sequences of XEC from high to low. Line, posterior mean, ribbon, 95% credible interval. **(E)** Lentivirus-based pseudovirus assay. HOS-ACE2/TMPRSS2 cells were infected with pseudoviruses bearing each S protein of KP.3 and KP.3.1.1. The amount of input virus was normalized to the amount of HIV-1 p24 capsid protein. The percentage infectivity of KP.3.1.1 is compared to that of KP.3. The horizontal dash line indicates the mean value of the percentage infectivity of KP.3. Assays were performed in quadruplicate, and a representative result of four independent assays is shown. The presented data is expressed as the average ± SD. Each dot indicates the result of an individual replicate. Statistically significant differences versus KP.3 is determined by two-sided Student’s *t* tests and statistically significant difference (*P* < 0.05) versus KP.3 is indicated with red asterisk. **(F)** Neutralization assay. Assays were performed with pseudoviruses harboring the S proteins of KP.3, KP.3.1.1, KP.3+T22N, KP.3+F59S and XEC. The following convalescent sera were used: sera from fully vaccinated individuals who had been infected with XBB.1.5 (one 2-dose vaccinated, three 3-dose vaccinated, five 4-dose vaccinated, three 5-dose vaccinated and one 6-dose vaccinated; time interval between the last vaccination and infection, 44-691 days; 15-46 days after testing. n=13 in total; average age: 44.1 years, range: 15-74 years, 30.8% male), individuals who had been infected with JN.1 (one 2-dose vaccinated, two 3-dose vaccinated, two 7-dose vaccinated and seven unknown vaccine history; time interval between the last vaccination and infection, 34–958 days; 13–46 days after testing. n=12 in total; average age: 69.3 years, range: 31–94 years, 41.7% male) and fully vaccinated individuals who had been infected with KP.3.3 (five 3-dose vaccinated, four 4-dose vaccinated, five 5-dose vaccinated and one 6-dose vaccinated; time interval between the last vaccination and infection, 208– 929 days; 13–45 days after testing. n=15 in total; average age: 48.1 years, range: 28–87 years, 46.7% male). Assays for each serum sample were performed in quadruplicate to determine the 50% neutralization titer (NT_50_). Each dot represents one NT_50_ value, and the median and 95% confidence interval are shown. The number in parenthesis indicates the geometric mean of NT_50_ values. The horizontal dash line indicates a detection limit (40-fold) and the number of serum donors with the NT_50_ values below the detection limit is shown in the figure (under the bars and dots of each variant). Neutralization titers below the detection limit were calculated as a titer of 40. Statistically significant differences versus KP.3 and KP.3.1.1 were determined by two-sided Wilcoxon signed-rank tests. The fold changes of NT_50_ versus KP.3 and KP.3.1.1 are calculated as the average of ratio of inverted NT_50_ obtained from each individual. The fold changes versus KP.3 are indicated with “X” followed by fold changes versus KP.3.1.1. Black and red numbers indicate fold changes versus KP.3 and KP.3.1.1 with statistically significant differences. Gray numbers indicate nonsignificant differences.

Thereafter, XEC, a recombinant lineage of KS.1.1 (JN.13.1.1.1) and KP.3.3 (JN.1.11.1.3.3), was first identified in Germany on August 7, 2024. XEC acquired two S substitutions, S:T22N and S:F59S, compared with KP.3 through recombination, with a breakpoint at genomic position 21,738-22,599 (**Figures 1A and 1B**). We estimated the relative effective reproduction number (R_e_) of XEC using a Bayesian multinomial logistic model^5^ based on genome surveillance data from the USA, the United Kingdom, France, Canada, and Germany, where this variant has spread as of August 2024 (**Figures 1C and 1D; Table S3**). In the USA, the R_e_ of XEC is 1.13-fold higher than that of KP.3.1.1 (**Figure 1C**). Additionally, the other countries under investigation herein showed higher R_e_ for XEC. These results suggest that XEC has the potential to outcompete the other major SARS-CoV-2 lineages including KP.3.1.1.^4^

We then assessed the virological properties of XEC using pseudoviruses. Pseudovirus infection assay showed that the infectivity of KP.3.1.1 and XEC was significantly higher than that of KP.3 (**Figure 1E**). Although S:T22N did not affect the infectivity of the pseudovirus based on KP.3, S:F59S significantly increased it (**Figure 1E**). Neutralization assay was performed using three types of human sera: convalescent sera after breakthrough infection (BTI) with XBB.1.5 or KP.3.3, and convalescent sera after JN.1 infection. In all serum groups, XEC as well as KP.3.1.1 showed immune resistance when compared to KP.3 with statistically significant differences (**Figure 1F**). In the cases of XBB.1.5 BTI sera and JN.1 infection sera, the 50% neutralization titers (NT_50_s) of XEC and KP.3.1.1 were comparable (**Figure 1F**). However, we revealed that the NT_50_ of XEC was significantly (1.3-fold) lower than that of KP.3.1.1 (**Figure 1F**). Moreover, both S:T22N and S:F59S significantly (1.5-fold and 1.6-fold) increased the resistance to KP.3.3 BTI sera (**Figure 1F**).

Altogether, here we showed that XEC exhibited higher pseudovirus infectivity and higher immune evasion than KP.3. Particularly, XEC exhibited more robust immune resistance to KP.3.3 BTI sera than KP.3.1.1. Our data suggest that the higher R_e_ of XEC than KP.3.1.1 is attributed to this property and XEC will be a predominant SARS-CoV-2 variant in the world in the near future.

