## Supplementary Appendix for "Virological characteristics of the SARS-CoV-2 XEC variant"

**Table of Contents**

**Contents** Page


| Contents | Page |
|---|---|
| **Materials and Methods** | 2-5 |
| Ethics statement | |
| Human serum collection | |
| Epidemic dynamics, mutation frequency and sequence polymorphism analysis | |
| Plasmid construction | |
| Cell culture | |
| Pseudovirus preparation | |
| Neutralization assay | |
| Data availability | |
| **Table S1.** Human infection sera used in this study | 6 |
| **Table S2.** Primers used in this study | 7 |
| **Table S3.** Estimated relative $R_e$ and epidemic dynamics modeling parameters of representative SARS-CoV-2 Omicron sublineages spreading in the USA, the United Kingdom, France, Canada, and Germany from January 1, 2024 to September 19, 2024 | 8-18 |
| **Consortia** | 19 |
| **Acknowledgments** | 20 |
| **Supplemental References** | 21 |

### Materials and Methods

#### Ethics statement

All protocols involving specimens from human subjects recruited at Kumamoto University, Interpark Kuramochi Clinic and Keio University were reviewed and approved by the Institutional Review Boards of The Institute of Medical Science, The University of Tokyo (approval IDs: 2021-1-0416 and 2022-29-0915), Kumamoto University (approval ID: 461), and Keio University (approval ID: 20200059), respectively. All human subjects provided written informed consent. All protocols for the use of human specimens were reviewed and approved by the Institutional Review Boards of The Institute of Medical Science, The University of Tokyo (approval IDs: 2021-1-0416, 2021-18-0617 and 2022-29-0915).

#### Human serum collection

Convalescent sera were collected from fully vaccinated individuals who had been infected with XBB.1.5 (one 2-dose vaccinated, three 3-dose vaccinated, five 4-dose vaccinated, three 5-dose vaccinated and one 6-dose vaccinated; time interval between the last vaccination and infection, 44-691 days; 15-46 days after testing. n=13 in total; average age: 44.1 years, range: 15-74 years, 30.8% male), individuals who had been infected with JN.1 (one 2-dose vaccinated, two 3-dose vaccinated, two 7-dose vaccinated and seven unknown vaccine history; time interval between the last vaccination and infection, 34–958 days; 13–46 days after testing. n=12 in total; average age: 69.3 years, range: 31–94 years, 41.7% male) and fully vaccinated individuals who had been infected with KP.3.3 (five 3-dose vaccinated, four 4-dose vaccinated, five 5-dose vaccinated and one 6-dose vaccinated; time interval between the last vaccination and infection, 208–929 days; 13–45 days after testing. n=15 in total; average age: 48.1 years, range: 28–87 years, 46.7% male). Sera were inactivated at 56°C for 30 minutes and stored at –80°C until use. The details of the convalescent sera are summarized in **Table S1**.

The infected SARS-CoV-2 variant was identified as previously described.[1-4] Briefly, the infected variant was determined by quantitative reverse-transcription polymerase chain reaction targeting of interests. The primers used for variant typing are listed in **Table S2**.

#### Epidemic dynamics, mutation frequency and sequence polymorphism analysis

In this study, we analyzed the viral genomic surveillance data stored in the GISAID database (https://www.gisaid.org; downloaded on October 9, 2024). We used the data of SARS-CoV-2 collected from January 1, 2024 to October 1, 2024 in this analysis. We excluded any data that i) lacks collection date and PANGO lineage information; ii) was retrieved from non-human animals; iii) was sampled by quarantine; iv) was sampled from the original passage; v) whose genomic sequence is not longer than 28,000 base pairs; and vi) contains >2% of unknown (N) nucleotide sequences. In the downstream analysis, we only used sequences for PANGO lineages with >40 sequences in each country in the dataset. We modeled the epidemic dynamics of variants of interest in the USA, the United Kingdom, France, Canada, and Germany where >100 genomic sequences of XEC were detected. The daily frequency of each viral lineage was counted. Then, epidemic dynamics and $R_e$ values for each viral lineage were subsequently estimated according to the Bayesian multinomial logistic model, as described in our previous study.[1] Briefly, we estimated the logistic slope parameter $\beta_l$ for each lineage and then calculated a relative $R_e$ for each lineage ($r_l$) as $r_l = \exp(\gamma\beta_l)$ where $\gamma$ is the average viral generation time (2.1 days) (http://sonorouschocolate.com/covid19/index.php?title=Estimating_Generation_Time_Of_Omicron). For parameter estimation, the intercept and slope parameters of KP.3.1.1 were fixed at 0. The relative $R_e$ of KP.3.1.1 was fixed at 1, and that of other lineage was estimated with respect to that of KP.3.1.1. Parameter estimation was performed by using the Markov chain Monte Carlo (MCMC) approach implemented in CmdStan v2.34.1 (https://mc-stan.org) accessed through the CmdStanr v0.6.1 R interface (https://mc-stan.org/cmdstanr/). Four independent 5,000-step MCMC chains were run including 1,000-step warmup iterations. We confirmed that an estimated $\hat{R}$ convergence diagnostic value is <1.05 and bulk and tail effective sampling sizes are >200, indicating that all runs were successfully convergent. Information on the estimated parameters of all variants is summarized in **Table S3**. Only JN.1, KP.2, KP.3, LB.1, KP.2.3, KP.3.1.1, and XEC variants are included in the **Figures 1C and 1D.** KP.3 and LB.1 are excluded from dataset of Germany due to lack of data in the **Figures 1C and 1D.**

Mutation frequency of each lineage was calculated by dividing the number of sequences harboring the substitution of interest with the total number of sequences in each lineage.

For sequence polymorphism analysis, the genomic sequence of XEC and its parental lineage, KS.1.1 and KP.3.3 were examined. We only used genomic sequences of variants collected in the USA and generated a consensus sequence for each lineage by retaining sites that were conserved in over 70% of the entire alignment. Nucleotide differences between the consensus sequences of the XEC, KS.1.1, and KP.3.3 lineages were visualized using snipit (https://github.com/aineniamh/snipit).

**Plasmid construction**

Plasmids expressing the SARS-CoV-2 spike proteins of KP.3 and KP.3.1.1 were prepared in our previous studies.[5-8] Plasmids expressing the spike protein of XEC and its derivatives were generated by site-directed overlap extension PCR using pC-SARS2-S KP.3 as the template and the primers listed in **Table S2**. The resulting PCR fragment was subcloned into the KpnI-NotI site of the pCAGGS vector[9] using In-Fusion HD Cloning Kit (Takara, Cat# Z9650N). Nucleotide sequences were determined by DNA sequencing services (Eurofins), and the sequence data were analyzed by SnapGene software v6.1.1 (www.snapgene.com).

**Cell culture**

The Lenti-X 293T cells (Takara, Cat# 632180) and HOS-ACE2/TMPRSS2 cells (kindly provided by Dr. Kenzo Tokunaga), a derivative of HOS cells (a human osteosarcoma cell line; ATCC CRL-1543) stably expressing human ACE2 and TMPRSS2,[10,11] were maintained in Dulbecco's modified Eagle's medium (DMEM) (high glucose) (Wako, Cat# 044- 29765) containing 10% fetal bovine serum (Sigma-Aldrich Cat# 172012-500ML), 100 units penicillin and 100 ug/ml streptomycin (Sigma-Aldrich, Cat# P4333-100ML).

**Pseudovirus preparation**

Pseudoviruses were prepared as previously described.[5,7,8,12] Briefly, lentivirus (HIV-1)-based, luciferase-expressing reporter viruses were pseudotyped with the SARS-CoV-2 S. One prior day of transfection, the LentiX-293T cells were seeded at a density of $2 \times 10^6$ cells. The LentiX-293T cells were cotransfected with 1 µg psPAX2-IN/HiBiT (a packaging plasmid encoding the HiBiT-tag-fused integrase[10]), 1 µg pWPI-Luc2 (a reporter plasmid encoding a firefly luciferase gene[13]) and 500 ng plasmids expressing parental S or its derivatives using TransIT-293 transfection reagent (Mirus, Cat# MIR2704) according to the manufacturer's protocol. Two days post transfection, the culture supernatants were harvested and filtrated. The amount of produced pseudovirus particles was quantified by the HiBiT assay using Nano Glo HiBiT lytic detection system (Promega, Cat# N3040) as previously described. [13] In this system, HiBiT peptide is produced with HIV-1 integrase and forms NanoLuc luciferase with LgBiT, which is supplemented with substrates. In each pseudovirus particle, the detected HiBiT value is correlated with the amount of the pseudovirus capsid protein, HIV-1 p24 protein. [13] Therefore, we calculated the amount of HIV-1 p24 capsid protein based on the HiBiT value measured, according to the previous paper. [13] To measure viral infectivity, the same amount of pseudovirus normalized with the HIV-1 p24 capsid protein was inoculated into HOS-ACE2/TMPRSS2 cells. At two days postinfection, the infected cells were lysed with a Bright-Glo luciferase assay system (Promega, Cat# E2620), and the luminescent signal produced by firefly luciferase reaction was measured using a GloMax explorer multimode microplate reader 3500 (Promega). The pseudoviruses were stored at –80°C until use.

**Neutralization assay**

Neutralization assays were performed previously described[5-8] with some modifications. The assays were mainly conducted by a semi-automated high-throughput method using Fluent780 (Tecan). [14,152,3] The SARS-CoV-2 spike pseudoviruses (counting ~100,000 relative light units) and serially diluted (40-fold to 29,160-fold dilution at the final concentration) heat-inactivated sera were manually prepared in a 2-ml 96-well plate (Greiner, Cat# 780271) and in 96-well microplates (ThermoFisher Scientific, Cat# 168136), respectively. The pseudoviruses were dispensed and mixed with the sera in 384-well plates (ThermoFisher Scientific, Cat# 164610) on Fluent780 (Tecan). Pseudoviruses without sera were included as controls. After incubation at 37°C for 1 hour, HOS-ACE2/TMPRSS2 cells (3,000 cells/30 μl) were added to the 20 μl mixture of pseudovirus and serum in the 384-well white plate on the device. Two days post infection, the infected cells were lysed with a Bright-Glo luciferase assay system (Promega, Cat# E2620) on Fluent780 (Tecan), and the luminescent signal was measured and processed using an Infinite200 and a Magellan (Tecan). The assay of each serum sample was performed in quadruplicate, and the 50% neutralization titer ($NT_{50}$) was calculated using Prism 9 (GraphPad Software).

### Data availability

The GISAID datasets used in this study are available from the GISAID database (https://www.gisaid.org; EPI-SET-ID: EPI_SET_241011ok, EPI_SET_241009gk and EPI_SET_241010ea). The supplemental tables for the GISAID datasets are available in the GitHub repository (https://github.com/TheSatoLab/XEC_short).

### Consortia

#### The Genotype to Phenotype Japan (G2P-Japan) Consortium

**The Institute of Medical Science, The University of Tokyo, Japan**
Naoko Misawa, Arnon Plianchaisuk, Ziyi Guo, Alfredo Hinay Jr., Kaoru Usui, Wilaiporn Saikruang, Spyridon Lytras, Daichi Yamasoba, Luca Nishimura, Shigeru Fujita, Jarel Elgin M. Tolentino, Lin Pan, Wenye Li, Maximilian Stanley Yo, Mai Suganami, Mika Chiba, Kyoko Yasuda, Keiko Iida, Adam P. Strange, Naomi Ohsumi, Shiho Tanaka, Eiko Ogawa, Tsuki Fukuda, Daniel Arnold

**Hokkaido University, Japan**
Takasuke Fukuhara, Tomokazu Tamura, Rigel Suzuki, Saori Suzuki, Shuhei Tsujino, Hayato Ito, Hirofumi Sawa, Naganori Nao, Keita Matsuno, Keita Mizuma, Jingshu Li, Izumi Kida, Yume Mimura, Yuma Ohari, Shinya Tanaka, Masumi Tsuda, Lei Wang, Yoshikata Oda, Zannatul Ferdous, Kenji Shishido, Hiromi Mohri, Miki Iida

**Tokyo Metropolitan Institute of Public Health**
Isao Yoshida

**Tokai University, Japan**
So Nakagawa

**Kyoto University, Japan**
Kotaro Shirakawa, Akifumi Takaori-Kondo, Kazuo Takayama, Rina Hashimoto, Sayaka Deguchi, Yukio Watanabe, Yoshitaka Nakata, Hiroki Futatsusako, Ayaka Sakamoto, Naoko Yasuhara, Takao Hashiguchi, Tateki Suzuki, Kanako Kimura, Jiei Sasaki, Yukari Nakajima, Hisano Yajima

**Hiroshima University, Japan**
Takashi Irie, Ryoko Kawabata

**Kyushu University, Japan**
Kaori Tabata

**Kumamoto University, Japan**
Hesham Nasser, Ryo Shimizu, Michael Jonathan, Yuka Mugita, Otowa Takahashi, Takamasa Ueno, Chihiro Motozono, Mako Toyoda

**University of Miyazaki, Japan**
Akatsuki Saito, Anon Kosaka, Miki Kawano, Natsumi Matsubara, Tomoko Nishiuchi

**Charles University, Czechia**
Jiri Zahradnik, Prokopios, Andrikopoulos, Miguel Padilla-Blanco, Aditi Konar, Ruojin Tuan

**Table S1. Human infection sera used in t**

| SARS-CoV-2 infected | Donor ID | Sex | Age | Number of vaccination | Date of 1st vaccination (YYYY-MM-DD) | Date of 2nd vaccination (YYYY-MM-DD) | Date of 3rd vaccination (YYYY-MM-DD) | Date of 4th vaccination (YYYY-MM-DD) | Date of 5th vaccination (YYYY-MM-DD) | Date of 6th vaccination (YYYY-MM-DD) | Date of 7th vaccination (YYYY-MM-DD) | Date of test (YYYY-MM-DD) | Date of sampling (YYYY-MM-DD) | Prior infection? |
|---|---|---|---|---|---|---|---|---|---|---|---|---|---|---|
| XBB.1.5 | 37306 | Female | 53 | 5 | 2021-04-27 (P) | 2021-05-18 (P) | 2022-02-01 (P) | 2022-07-30 (M) | 2022-12-17 (P) | | | 2023-07-20 | 2023-08-08 | No |
| XBB.1.5 | 37072 | Female | 15 | 3 | 2021-09-25 (P) | 2021-10-18 (P) | 2022-05-02 (P) | | | | | 2023-07-11 | 2023-08-11 | No |
| XBB.1.5 | 37071 | Female | 48 | 3 | 2021-10-01 (P) | 2021-11-01 (P) | 2022-05-06 (P) | | | | | 2023-07-15 | 2023-08-11 | No |
| XBB.1.5 | 36845 | Male | 29 | 3 | 2021-09-01 (M) | 2021-09-29 (M) | 2022-05-27 (M) | | | | | 2023-06-26 | 2023-08-11 | No |
| XBB.1.5 | 37229 | Female | 74 | 5 | 2021-06-24 (P) | 2021-07-15 (P) | 2022-02-16 (M) | 2022-07-20 (M) | 2023-03-25 (M) | | | 2023-07-17 | 2023-08-01 | No |
| XBB.1.5 | 37998 | Female | 65 | 6 | 2021-08-04 (P) | 2021-08-30 (P) | 2022-03-19 (M) | 2022-09-02 (P) | 2022-12-24 (P) | 2023-06-27 (PBA4/5) | | 2023-08-10 | 2023-09-02 | No |
| XBB.1.5 | 37798 | Female | 62 | 5 | 2021-03-17 (P) | 2021-04-09 (P) | 2021-12-23 (P) | 2022-07-28 (P) | 2023-06-17 (P) | | | 2023-08-03 | 2023-08-20 | No |
| XBB.1.5 | 38019 | Female | 18 | 4 | 2021-09-07 (P) | 2021-10-07 (P) | 2022-04-28 (P) | 2022-12-27 (P) | | | | 2023-08-10 | 2023-09-04 | No |
| XBB.1.5 | 38871 | Male | 46 | 4 | 2021-09-12 (P) | 2021-10-03 (P) | 2022-04-08 (M) | 2022-11-05 (M) | | | | 2023-08-22 | 2023-09-10 | No |
| XBB.1.5 | 38880 | Male | 51 | 4 | 2021-10-07 (P) | 2021-10-28 (P) | 2022-05-13 (M) | 2022-11-09 (PBA4/5) | | | | 2023-08-22 | 2023-09-16 | No |
| XBB.1.5 | 39296 | Male | 48 | 4 | 2021-07-15 (M) | 2021-08-23 (M) | 2022-03-24 (M) | 2022-12-25 (M) | | | | 2023-09-01 | 2023-09-23 | No |
| XBB.1.5 | 39502 | Female | 43 | 4 | 2021-07-31 (P) | 2021-08-23 (P) | 2022-03-15 (P) | 2022-11-01 (P) | | | | 2023-09-05 | 2023-09-26 | No |
| XBB.1.5 | 37463 | Female | 21 | 2 | 2021-08 (M) | 2021-09 (M) | | | | | | 2023-07-24 | 2023-08-12 | No |
| JN.1 | 3315 | Female | 75 | 7 | NA | NA | NA | NA | NA | NA | 2023-11-07 | 2023-12-11 | 2024-01-15 | No |
| JN.1 | 3323 | Female | 73 | 3 | 2021 | 2021 | 2021-05 | | | | | 2023-12-15 | 2023-12-28 | Yes |
| JN.1 | 3325 | Female | 73 | 2 | 2021-06 | 2021-07 | | | | | | 2023-12-16 | 2024-01-16 | NA |
| JN.1 | 3338 | Male | 31 | 3 | 2021-06 | 2021-07 | 2022-02-02 | | | | | 2023-12-26 | 2024-02-10 | Yes |
| JN.1 | 3355 | Male | 54 | 7 | NA | NA | NA | NA | NA | NA | NA | 2024-01-03 | 2024-02-15 | NA |
| JN.1 | 3316 | Female | 52 | NA | ? | | | | | | | 2023-12-11 | 2024-01-06 | NA |
| JN.1 | 3320 | Female | 80 | NA | ? | | | | | | | 2023-12-13 | 2024-01-12 | NA |
| JN.1 | 3329 | Male | 94 | NA | ? | | | | | | | 2023-12-18 | 2024-01-23 | NA |
| JN.1 | 3337 | Male | 72 | NA | ? | | | | | | | 2023-12-26 | 2024-01-19 | NA |
| JN.1 | 3362 | Male | 74 | NA | ? | | | | | | | 2024-01-06 | 2024-01-19 | NA |
| JN.1 | 3367 | Female | 70 | NA | ? | | | | | | | 2024-01-08 | 2024-01-26 | NA |
| JN.1 | 3400 | Female | 84 | NA | ? | | | | | | | 2024-01-24 | 2024-02-09 | NA |
| KP.3.3 | NS 02 | Female | 40 | 4 | 2021-08-08 (M) | 2021-09-04 (M) | 2022-04-22 (P) | 2022-10-13 (MBA1/2) | | | | 2024-07-23 | 2024-08-06 | Yes |
| KP.3.3 | NS 03 | Male | 40 | 5 | 2021-07-24 (P) | 2021-08-16 (P) | 2022-04-16 (P) | 2022-10-15 (P) | 2023-11-22 (P) | | | 2024-07-24 | 2024-08-06 | Yes |
| KP.3.3 | NS 04 | Female | 63 | 6 | 2021-05-26 (P) | 2021-06-16 (P) | 2022-02-12 (P) | 2022-07-25 (P) | 2022-11-25 (P) | 2023-12-05 (P) | | 2024-07-20 | 2024-08-06 | Yes |
| KP.3.3 | NS 05 | Male | 35 | 3 | 2021-04-28 (P) | 2021-05-19 (P) | 2022-01-06 (P) | | | | | 2024-07-23 | 2024-08-09 | Yes |
| KP.3.3 | NS 06 | Female | 37 | 3 | 2021-03-05 (P) | 2021-05-21 (P) | 2022-02-14 (P) | | | | | 2024-07-21 | 2024-08-08 | Yes |
| KP.3.3 | NS 07 | Female | 57 | 3 | 2021-08-20 (M) | 2021-09-17 (M) | 2022-05-18 (M) | | | | | 2024-07-22 | 2024-08-06 | Yes |
| KP.3.3 | NS 08 | Female | 38 | 3 | 2021-03-03 (A) | 2021-05-19 (A) | 2022-04-04 (P) | | | | | 2024-07-21 | 2024-08-08 | Yes |
| KP.3.3 | NS 10 | Male | 28 | 4 | 2021-08-23 (M) | 2021-09-22 (M) | 2022-03-26 (P) | 2023-01-07 (P) | | | | 2024-07-22 | 2024-08-06 | Yes |
| KP.3.3 | NS 11 | Male | 41 | 5 | 2021-07-12 (M) | 2021-08-06 (M) | 2022-03-18 (M) | 2022-12-09 (P) | 2023-12-28 (P) | | | 2024-07-23 | 2024-08-07 | Yes |
| KP.3.3 | 3403 | Male | 31 | 5 | NA | NA | NA | NA | NA | | | 2024-06-30 | 2024-07-15 | NA |
| KP.3.3 | 3407 | Female | 72 | 5 | NA | NA | NA | NA | NA | | | 2024-07-01 | 2024-08-05 | NA |
| KP.3.3 | 3408 | Male | 87 | 5 | NA | NA | NA | NA | NA | | | 2024-06-27 | 2024-07-16 | NA |
| KP.3.3 | 3415 | Female | 44 | 4 | NA | NA | NA | NA | | | | 2024-07-02 | 2024-07-24 | NA |
| KP.3.3 | 3431 | Female | 74 | 4 | NA | NA | NA | NA | | | | 2024-07-08 | 2024-08-13 | NA |
| KP.3.3 | 3463 | Male | 34 | 3 | NA | NA | NA | | | | | 2024-07-15 | 2024-08-29 | NA |

NA, not applicable.

A, Astrazeneca; P, Pfizer-BioNTech; M, Moderna

BA1/2, BA.1/2 bivalent vaccine; BA4/5, BA.4/5 bivalent vaccine

**Table S2. Primers used in this study**

| Target region | Primer name | Primer sequence (5'-to-3') | Purpose |
|---|---|---|---|
| ORF1a:T4175I | ORF1a_T4175I-F | GCACTGATGACAATGCGTTAGC | Variant typing |
| | ORF1a_T4175I-R | CCTGTAAATCGGATAACAGTGCAAG | Variant typing |
| | ORF1a_T4175c | FAM-AACACAAcAAAGGGAG-MGB | Variant typing |
| | ORF1a_I4175t | VIC-ACAACACAAtAAAGGGAGG-MGB | Variant typing |
| S:S31del | S_S51 | FAM-ATACACTAATtctTTCACACGTG-MGB | Variant typing |
| | S_S31del | VIC-CATACACTAAT---TTCACACGTG-MGB | Variant typing |
| | S_S31del-F | GCCACTAGTCTCTAGTCAGTGTGTTAATC | Variant typing |
| | S_S31del-R | GAGGATCTGAAAACTTTGTCAGGG | Variant typing |
| S:Q183H | S_Q183g | FAM-GGAAAACAgGGTAATT-MGB | Variant typing |
| | S_H183t | VIC-AGGAAAACAtGGTAATTT-MGB | Variant typing |
| | S_Q183H-F | CTCAGCCTTTTCTTATGGACCTTG | Variant typing |
| | S_Q183H-R | CCATCAATATTCTTAAACACAAATTCCC | Variant typing |
| S:Q183E | S_Q183E-F | GTCTCTCAGCCTTTTCTTATGGACC | Variant typing |
| | S_Q183E-R | CCATCAATATTCTTAAACACAAATTCCC | Variant typing |
| | S_Q183t | FAM-GAAGGAAAAcAGGGTAAT-MGB | Variant typing |
| | S_E183c | VIC-GAAGGAAAAgAGGGTAAT-MGB | Variant typing |
| S:R346T | S_R346g | FAM-CACCAgATTTGC-MGB | Variant typing |
| | S_T346c | VIC-CCACCAcATTTGC-MGB | Variant typing |
| | S_R346T-F | AAACTTGTGCCCTTTTGATGAAG | Variant typing |
| | S_R346T-R | CTGATTCTCTTCCTGTTCCAAGC | Variant typing |
| S: L455S | S_L455S-F | CTAACAAGCTTGATTCTAAGCATAGTGG | Variant typing |
| | S_L455S-R | GTTGAAATATCTCTCTCAAAAGGTTTGAG | Variant typing |
| | S_L455 | FAM-CTGGTATAGATtGTTTAGGAAG-MGB | Variant typing |
| | S_S455 | VIC-TGGTATAGATcGTTTAGGAAG-MGB | Variant typing |
| S: L455F | S_L455F_F456L-Fw | GCTTGGAATTCTAACAAGCTTGATT | Variant typing |
| | S_L455F_F456L-Rv | TCAGTTGAAATATCTCTCTCAAAAGGTT | Variant typing |
| | S_L455g-L456a | FAM-TAGATTgTTaAGGAAGTCTA-MGB | Variant typing |
| | S_F455t-L456a | VIC-TAGATTtTTaAGGAAGTCTAAGC-MGB | Variant typing |
| S_L455S_F456L | S_S455c-F456t | FAM GTATAGAT c G t TTAGGAAGT MGB | Variant typing |
| | S_S455c-L456c | VIC-GTATAGATcGcTTAGGAAG-MGB | Variant typing |
| | S_L455S_F456L-F | AACAAGCTTGATTCTAAGCATAGTGG | Variant typing |
| | S_L455S_F456L-R | GCCTGATAGATTTCAGTTGAAATATCTC | Variant typing |
| S: F456L | S_F456L-F | ACAAGCTTGATTCTAAGCCTAGTGG | Variant typing |
| | S_F456L-R | GTTGAAATATCTCTCTCAAAAGGTTTGA | Variant typing |
| | S_F456t | FAM-ATAGATTGTTtAGGAAGTCT-MGB | Variant typing |
| | S_L456a | VIC-ATAGATTGTTaAGGAAGTCT-MGB | Variant typing |
| S: V483del | S_V483del-F | CAACTGAAATCTATCAGGCCGG | Variant typing |
| | S_V483del-R | GGGTCGGAAACCATATGATTGTAA | Variant typing |
| | S_V483 | FAM-TTGTAATGGTGTTGCAGGT-MGB | Variant typing |
| | S_V483del | VIC-CTTGTAAAGGTAAAGGTCCTA-MGB | Variant typing |
| S: S486P | S_S486P-F | CAGGCCGGTAACAAACCTTG | Variant typing |
| | S_S486P-R | CGGAAACCATATGATTGTAAAGGAG | Variant typing |
| | S_S486t | FAM-TGCAGGTtCTAATTGT-MGB | Variant typing |
| | S_P486c | VIC-GCAGGTcCTAATTG-MGB | Variant typing |
| S protein | Omicron universal Fw | cactatagggcgaattgggtaccatgtttgtgttcctggt | Expression plasmid preparation |
| | BA.2 WT Rv | agctccaccgcggtggcggccgctcaggtgtagtgcagtttca | Expression plasmid preparation |
| | KP3_T22N-Fw | accACCAATcaaAGCtacaccaactccttc | Expression plasmid preparation |
| | KP3_T22N-Rv | GCTttgATTGGTggtGATcaggttGAACAG | Expression plasmid preparation |
| | KP3_F59S-Fw | tgccattcTCCagcaatgtgacctggttcc | Expression plasmid preparation |
| | KP3_F59S-Rv | attgctGGAgaatggcaggaacaggtcctg | Expression plasmid preparation |

**Table S3. Estimated relative Re and epidemic dynamics modeling parameters of the representative SARS-CoV-2 Omicron sublineages spreading in USA, Germany, United Kingdom, Canada, and France from Jan 1, 2024 to September 19, 2024**

| | | Relative $R_e$ (posterior values) | | | | | |
|---|---|---|---|---|---|---|---|
| PANGO lineage | Country | Mean | $2.5^{th}$ percentile | $97.5^{th}$ percentile | $\hat{R}$ | Bulk effective sample size | Tail effective sample size |
| XEC | USA | 1.129 | 1.108 | 1.152 | 1.000 | 19170.1 | 11311.2 |
| MC.1 | USA | 1.059 | 1.041 | 1.077 | 1.001 | 16651.1 | 11880.5 |
| MB.1.1 | USA | 1.021 | 1.001 | 1.043 | 1.000 | 18001.1 | 11135.0 |
| KP.3.3.2 | USA | 1.002 | 0.986 | 1.020 | 1.000 | 15866.6 | 11433.6 |
| JN.1.18.6 | USA | 0.998 | 0.977 | 1.021 | 1.001 | 17328.4 | 11521.2 |
| KP.2.3.4 | USA | 0.989 | 0.969 | 1.010 | 1.000 | 13340.2 | 11436.4 |
| LB.1.1 | USA | 0.981 | 0.963 | 0.999 | 1.001 | 17687.9 | 11452.3 |
| KP.3.3.1 | USA | 0.980 | 0.968 | 0.993 | 1.000 | 11517.6 | 11468.6 |
| XDY | USA | 0.978 | 0.962 | 0.994 | 1.000 | 14378.5 | 11367.3 |
| LY.1 | USA | 0.975 | 0.965 | 0.984 | 1.000 | 6035.7 | 11984.5 |
| MK.1 | USA | 0.971 | 0.956 | 0.988 | 1.000 | 14763.6 | 11633.1 |
| KP.3.2.1 | USA | 0.970 | 0.951 | 0.990 | 1.000 | 16228.6 | 11220.5 |
| KP.2.14 | USA | 0.970 | 0.957 | 0.983 | 1.000 | 11727.5 | 10684.1 |
| LB.1.4.1 | USA | 0.967 | 0.951 | 0.983 | 1.000 | 15078.6 | 11829.1 |
| LB.1.5 | USA | 0.966 | 0.948 | 0.984 | 1.001 | 16320.0 | 11463.8 |
| LB.1.7.1 | USA | 0.963 | 0.947 | 0.981 | 1.000 | 15700.8 | 11282.1 |
| LU.2 | USA | 0.963 | 0.944 | 0.984 | 1.000 | 15638.0 | 10918.8 |
| KP.2.6 | USA | 0.959 | 0.944 | 0.976 | 1.000 | 13179.9 | 10469.1 |
| LB.1.7.2 | USA | 0.958 | 0.942 | 0.975 | 1.000 | 15860.0 | 12343.0 |
| LB.1.2 | USA | 0.957 | 0.949 | 0.966 | 1.001 | 3919.3 | 9538.8 |
| KP.1.1.5 | USA | 0.957 | 0.950 | 0.964 | 1.001 | 2382.9 | 7792.1 |
| KS.1.1 | USA | 0.956 | 0.948 | 0.965 | 1.000 | 4604.2 | 9103.2 |
| LW.1 | USA | 0.956 | 0.941 | 0.971 | 1.000 | 12353.9 | 11726.6 |
| LB.1.2.1 | USA | 0.955 | 0.944 | 0.967 | 1.001 | 7808.7 | 11541.4 |
| LP.1 | USA | 0.954 | 0.949 | 0.960 | 1.001 | 1601.4 | 5344.7 |
| KP.3.5 | USA | 0.952 | 0.945 | 0.960 | 1.000 | 3543.2 | 9076.1 |
| KP.3.1.3 | USA | 0.951 | 0.935 | 0.968 | 1.000 | 16260.6 | 11536.8 |
| KP.3.1.6 | USA | 0.951 | 0.941 | 0.961 | 1.000 | 6972.0 | 12026.1 |
| MA.1 | USA | 0.948 | 0.936 | 0.961 | 1.001 | 7731.1 | 9729.1 |
| KP.3.4 | USA | 0.948 | 0.931 | 0.966 | 1.000 | 12267.4 | 11403.1 |
| LP.1.1 | USA | 0.948 | 0.931 | 0.966 | 1.000 | 14640.6 | 11758.5 |
| LB.1.3 | USA | 0.948 | 0.944 | 0.952 | 1.003 | 796.3 | 2368.0 |
| KP.2.3.2 | USA | 0.947 | 0.932 | 0.963 | 1.000 | 12447.9 | 11508.0 |
| KP.2.3 | USA | 0.946 | 0.943 | 0.949 | 1.006 | 402.9 | 1228.7 |
| KP.4.1.3 | USA | 0.946 | 0.934 | 0.959 | 1.000 | 9076.1 | 10592.6 |
| LB.1.7 | USA | 0.946 | 0.942 | 0.950 | 1.003 | 732.6 | 2553.1 |

| | | | | | | | |
|---|---|---|---|---|---|---|---|
| KP.2.3.3 | USA | 0.945 | 0.934 | 0.957 | 1.000 | 8416.9 | 11264.0 |
| KP.1.1.3 | USA | 0.944 | 0.940 | 0.948 | 1.004 | 833.3 | 3670.1 |
| LB.1.4 | USA | 0.942 | 0.934 | 0.951 | 1.001 | 3760.2 | 9070.8 |
| KP.2.15 | USA | 0.940 | 0.932 | 0.948 | 1.001 | 3414.7 | 8850.6 |
| KP.3.2.4 | USA | 0.939 | 0.928 | 0.951 | 1.001 | 6236.0 | 10936.7 |
| LF.3.1 | USA | 0.939 | 0.928 | 0.950 | 1.001 | 6060.8 | 11200.5 |
| LB.1.8 | USA | 0.938 | 0.928 | 0.949 | 1.001 | 5443.2 | 10145.3 |
| JN.1.7.8 | USA | 0.937 | 0.922 | 0.953 | 1.000 | 9492.5 | 11968.7 |
| KP.3.2.5 | USA | 0.937 | 0.925 | 0.950 | 1.000 | 9296.9 | 11209.8 |
| KP.3.3.3 | USA | 0.936 | 0.930 | 0.943 | 1.001 | 2412.7 | 7065.8 |
| KP.2.9 | USA | 0.936 | 0.929 | 0.942 | 1.001 | 2175.9 | 7509.0 |
| JN.1.11 | USA | 0.934 | 0.930 | 0.939 | 1.002 | 998.1 | 3699.4 |
| LD.1 | USA | 0.934 | 0.921 | 0.947 | 1.001 | 8863.4 | 10921.8 |
| KP.3.2.3 | USA | 0.933 | 0.929 | 0.938 | 1.003 | 856.0 | 3577.6 |
| LF.1.1.1 | USA | 0.931 | 0.921 | 0.943 | 1.000 | 7375.9 | 10809.9 |
| LQ.1 | USA | 0.928 | 0.912 | 0.945 | 1.000 | 12714.2 | 12431.9 |
| KS.1.3 | USA | 0.927 | 0.911 | 0.944 | 1.000 | 11085.2 | 10906.0 |
| KP.3.3 | USA | 0.925 | 0.922 | 0.929 | 1.005 | 455.8 | 1421.1 |
| ML.2 | USA | 0.925 | 0.916 | 0.934 | 1.001 | 3990.7 | 10896.1 |
| LF.3.1.1 | USA | 0.923 | 0.915 | 0.932 | 1.000 | 3840.6 | 9302.9 |
| JN.1.9.2 | USA | 0.922 | 0.911 | 0.933 | 1.000 | 7325.6 | 10862.8 |
| JN.1.50 | USA | 0.922 | 0.908 | 0.937 | 1.000 | 11415.9 | 10380.6 |
| KP.2.2 | USA | 0.922 | 0.918 | 0.925 | 1.004 | 575.6 | 1933.1 |
| KP.3.1 | USA | 0.921 | 0.918 | 0.925 | 1.005 | 493.3 | 1437.2 |
| LB.1 | USA | 0.919 | 0.915 | 0.922 | 1.004 | 487.2 | 1748.0 |
| KP.3 | USA | 0.918 | 0.915 | 0.922 | 1.004 | 593.0 | 1920.6 |
| KP.3.1.4 | USA | 0.917 | 0.911 | 0.923 | 1.001 | 1721.8 | 5241.1 |
| JN.1.16.3 | USA | 0.915 | 0.906 | 0.924 | 1.001 | 3811.3 | 9828.0 |
| LQ.1.1 | USA | 0.915 | 0.905 | 0.924 | 1.001 | 3691.4 | 9647.3 |
| KP.3.2 | USA | 0.912 | 0.907 | 0.916 | 1.002 | 847.3 | 2922.8 |
| KP.1.1.1 | USA | 0.909 | 0.905 | 0.913 | 1.003 | 680.3 | 2384.7 |
| JN.1.16.1 | USA | 0.908 | 0.904 | 0.911 | 1.006 | 465.0 | 1534.6 |
| KP.5 | USA | 0.907 | 0.896 | 0.918 | 1.000 | 6040.6 | 11076.5 |
| XDV.1 | USA | 0.904 | 0.899 | 0.910 | 1.002 | 1151.4 | 4921.7 |
| KP.2.16 | USA | 0.904 | 0.894 | 0.914 | 1.001 | 4083.4 | 10445.8 |
| KR.1.1 | USA | 0.903 | 0.890 | 0.915 | 1.000 | 8026.9 | 10837.0 |
| JN.1.7.4 | USA | 0.901 | 0.890 | 0.913 | 1.001 | 5519.1 | 11569.5 |
| JN.1.48.1 | USA | 0.901 | 0.891 | 0.911 | 1.001 | 4599.2 | 9408.7 |
| KP.4 | USA | 0.899 | 0.889 | 0.910 | 1.001 | 4605.3 | 10782.8 |
| KP.2 | USA | 0.899 | 0.896 | 0.902 | 1.007 | 331.1 | 1032.4 |

| | | | | | | | |
|---|---|---|---|---|---|---|---|
| JN.1.18.4 | USA | 0.898 | 0.886 | 0.910 | 1.001 | 5602.2 | 10271.7 |
| KS.1 | USA | 0.896 | 0.891 | 0.900 | 1.003 | 674.1 | 2470.9 |
| JN.1.18.3 | USA | 0.895 | 0.886 | 0.904 | 1.000 | 4256.9 | 8883.4 |
| JN.1.40 | USA | 0.895 | 0.887 | 0.903 | 1.001 | 2880.0 | 9819.6 |
| KP.2.1 | USA | 0.894 | 0.884 | 0.905 | 1.001 | 5056.0 | 10295.4 |
| KP.4.2 | USA | 0.894 | 0.885 | 0.903 | 1.001 | 2742.8 | 8320.0 |
| KP.1.1 | USA | 0.893 | 0.889 | 0.896 | 1.005 | 489.6 | 1593.8 |
| JN.1.16 | USA | 0.893 | 0.889 | 0.896 | 1.006 | 419.5 | 1369.8 |
| KP.4.1 | USA | 0.892 | 0.887 | 0.897 | 1.003 | 1011.2 | 3477.2 |
| KW.1.1 | USA | 0.891 | 0.886 | 0.897 | 1.002 | 1068.1 | 4097.6 |
| KP.1.2 | USA | 0.890 | 0.883 | 0.896 | 1.002 | 1671.0 | 5563.6 |
| JN.1.18.5 | USA | 0.889 | 0.879 | 0.899 | 1.000 | 4021.7 | 10015.6 |
| JN.1.11.1 | USA | 0.886 | 0.882 | 0.890 | 1.003 | 596.7 | 1598.8 |
| JN.1.18.2 | USA | 0.885 | 0.877 | 0.893 | 1.001 | 2658.6 | 6865.4 |
| JN.1.42.2 | USA | 0.885 | 0.876 | 0.894 | 1.001 | 3771.7 | 8885.7 |
| KR.1 | USA | 0.884 | 0.878 | 0.890 | 1.002 | 1271.2 | 5051.1 |
| JN.1.48 | USA | 0.884 | 0.874 | 0.894 | 1.001 | 3387.9 | 10014.0 |
| LA.1 | USA | 0.882 | 0.875 | 0.889 | 1.001 | 2127.9 | 6826.5 |
| LF.1 | USA | 0.879 | 0.870 | 0.889 | 1.001 | 3489.9 | 9635.9 |
| JN.1.10 | USA | 0.876 | 0.868 | 0.884 | 1.001 | 3296.9 | 7927.5 |
| KU.2 | USA | 0.875 | 0.866 | 0.884 | 1.000 | 3604.5 | 8753.2 |
| JN.1.37 | USA | 0.872 | 0.866 | 0.878 | 1.001 | 1454.3 | 4663.1 |
| XDK.1 | USA | 0.870 | 0.862 | 0.879 | 1.000 | 3222.9 | 7745.3 |
| JN.1.7.3 | USA | 0.868 | 0.859 | 0.877 | 1.001 | 2725.8 | 7643.3 |
| JN.1.8 | USA | 0.863 | 0.859 | 0.866 | 1.006 | 426.2 | 1292.5 |
| JN.1.1.6 | USA | 0.858 | 0.849 | 0.867 | 1.001 | 3196.0 | 8277.7 |
| XDQ | USA | 0.858 | 0.851 | 0.865 | 1.001 | 1945.8 | 5924.2 |
| JN.1.4.4 | USA | 0.856 | 0.852 | 0.859 | 1.006 | 476.3 | 1555.6 |
| JN.1.15 | USA | 0.855 | 0.848 | 0.861 | 1.002 | 1657.6 | 6400.2 |
| JN.1.18 | USA | 0.855 | 0.851 | 0.858 | 1.006 | 412.5 | 1199.1 |
| JN.1.26 | USA | 0.854 | 0.847 | 0.861 | 1.001 | 1875.8 | 7090.0 |
| JN.1.3 | USA | 0.853 | 0.847 | 0.859 | 1.002 | 1414.2 | 4117.2 |
| JN.1.20 | USA | 0.852 | 0.846 | 0.858 | 1.002 | 1437.9 | 5329.8 |
| JN.1.24.1 | USA | 0.852 | 0.843 | 0.860 | 1.001 | 3056.9 | 8261.5 |
| JN.1.59 | USA | 0.851 | 0.842 | 0.860 | 1.000 | 3239.4 | 7932.0 |
| KW.1 | USA | 0.851 | 0.842 | 0.860 | 1.001 | 3250.4 | 8495.2 |
| JN.1.56 | USA | 0.848 | 0.839 | 0.857 | 1.001 | 3720.2 | 8725.2 |
| KP.1 | USA | 0.848 | 0.842 | 0.854 | 1.002 | 1177.3 | 3964.3 |
| JN.1.1.3 | USA | 0.847 | 0.837 | 0.858 | 1.000 | 4335.1 | 10878.4 |
| JN.1.30 | USA | 0.847 | 0.839 | 0.856 | 1.000 | 3075.2 | 8571.7 |

| | | | | | | | |
|---|---|---|---|---|---|---|---|
| JN.1.34 | USA | 0.847 | 0.842 | 0.853 | 1.002 | 1173.6 | 3662.2 |
| KQ.1 | USA | 0.847 | 0.843 | 0.851 | 1.004 | 676.2 | 2261.2 |
| JN.1.4.8 | USA | 0.845 | 0.835 | 0.856 | 1.001 | 3999.5 | 8713.0 |
| JN.1.13.1 | USA | 0.845 | 0.841 | 0.848 | 1.006 | 380.6 | 1103.4 |
| JN.1.32 | USA | 0.844 | 0.840 | 0.848 | 1.005 | 493.5 | 1671.0 |
| JN.1.7 | USA | 0.843 | 0.841 | 0.846 | 1.008 | 291.1 | 783.3 |
| JN.1.63 | USA | 0.843 | 0.833 | 0.853 | 1.001 | 3536.6 | 8256.7 |
| JN.1.13 | USA | 0.842 | 0.831 | 0.853 | 1.001 | 5576.0 | 11014.8 |
| JN.1.52 | USA | 0.842 | 0.836 | 0.848 | 1.002 | 1354.0 | 5165.0 |
| JN.1.4.3 | USA | 0.841 | 0.836 | 0.847 | 1.002 | 1017.8 | 3858.2 |
| XDP.1 | USA | 0.841 | 0.835 | 0.847 | 1.002 | 1399.6 | 5271.0 |
| JN.1.7.2 | USA | 0.837 | 0.832 | 0.842 | 1.002 | 891.5 | 3197.1 |
| JN.1.4.6 | USA | 0.836 | 0.830 | 0.842 | 1.002 | 1214.8 | 4752.8 |
| JN.1.53 | USA | 0.832 | 0.821 | 0.842 | 1.001 | 4159.7 | 10664.9 |
| JN.1.60 | USA | 0.832 | 0.819 | 0.843 | 1.000 | 5536.5 | 10056.6 |
| JN.1.58 | USA | 0.831 | 0.820 | 0.841 | 1.001 | 4569.3 | 8263.5 |
| XDK | USA | 0.830 | 0.822 | 0.839 | 1.001 | 2939.6 | 8581.7 |
| JN.1.8.3 | USA | 0.827 | 0.815 | 0.838 | 1.001 | 5676.7 | 10156.3 |
| JN.1.8.1 | USA | 0.826 | 0.823 | 0.829 | 1.007 | 339.1 | 987.8 |
| JN.1.1.5 | USA | 0.825 | 0.815 | 0.835 | 1.000 | 4825.8 | 9396.3 |
| KV.2 | USA | 0.825 | 0.821 | 0.829 | 1.004 | 642.8 | 2426.8 |
| JN.1.7.5 | USA | 0.825 | 0.820 | 0.829 | 1.004 | 657.2 | 2833.2 |
| JN.1.1.1 | USA | 0.825 | 0.811 | 0.837 | 1.000 | 6146.6 | 11311.4 |
| XDP | USA | 0.823 | 0.818 | 0.828 | 1.003 | 806.1 | 2946.1 |
| BA.2 | USA | 0.821 | 0.813 | 0.830 | 1.001 | 3030.5 | 7473.5 |
| JN.1.14 | USA | 0.820 | 0.810 | 0.829 | 1.001 | 3757.8 | 8414.0 |
| JN.1.55 | USA | 0.814 | 0.802 | 0.826 | 1.000 | 5944.2 | 11216.0 |
| JN.1.43.1 | USA | 0.814 | 0.807 | 0.820 | 1.002 | 1617.3 | 5426.2 |
| JN.1.5 | USA | 0.809 | 0.801 | 0.816 | 1.001 | 2159.8 | 7101.3 |
| JN.1.39 | USA | 0.808 | 0.804 | 0.812 | 1.004 | 587.4 | 2233.4 |
| JN.1.21 | USA | 0.804 | 0.793 | 0.816 | 1.000 | 5952.5 | 10596.5 |
| JN.1.9 | USA | 0.804 | 0.800 | 0.809 | 1.003 | 833.5 | 3015.1 |
| JN.1.4.2 | USA | 0.804 | 0.799 | 0.809 | 1.003 | 854.6 | 3526.6 |
| JN.1.4.7 | USA | 0.801 | 0.792 | 0.810 | 1.000 | 3217.4 | 8152.9 |
| JN.1.42 | USA | 0.801 | 0.796 | 0.806 | 1.003 | 812.8 | 2619.1 |
| JN.1.29 | USA | 0.801 | 0.790 | 0.811 | 1.000 | 5450.8 | 9291.9 |
| JN.1.49 | USA | 0.800 | 0.784 | 0.815 | 1.000 | 9019.6 | 10984.9 |
| JN.1.4 | USA | 0.799 | 0.796 | 0.802 | 1.008 | 298.1 | 825.8 |
| JN.1.17 | USA | 0.798 | 0.789 | 0.807 | 1.001 | 3382.7 | 7214.5 |
| JN.1 | USA | 0.798 | 0.795 | 0.801 | 1.009 | 280.3 | 768.4 |

| | | | | | | | |
|---|---|---|---|---|---|---|---|
| JN.1.2 | USA | 0.797 | 0.792 | 0.803 | 1.002 | 1044.2 | 3969.5 |
| JN.1.38 | USA | 0.797 | 0.790 | 0.803 | 1.002 | 1643.5 | 4813.9 |
| JN.1.19 | USA | 0.796 | 0.789 | 0.803 | 1.001 | 2103.6 | 5217.0 |
| JN.1.22 | USA | 0.796 | 0.790 | 0.802 | 1.002 | 1337.4 | 4642.2 |
| JN.1.4.5 | USA | 0.795 | 0.792 | 0.799 | 1.005 | 450.1 | 1272.6 |
| JN.1.4.1 | USA | 0.794 | 0.777 | 0.811 | 1.000 | 10421.1 | 10762.4 |
| JN.1.64 | USA | 0.794 | 0.780 | 0.807 | 1.000 | 7198.6 | 10295.0 |
| JN.1.6 | USA | 0.793 | 0.787 | 0.799 | 1.002 | 1388.2 | 3969.8 |
| JN.1.27 | USA | 0.792 | 0.776 | 0.807 | 1.000 | 8525.3 | 11263.4 |
| JN.1.46 | USA | 0.792 | 0.786 | 0.798 | 1.002 | 1621.1 | 5090.4 |
| XDD | USA | 0.790 | 0.781 | 0.799 | 1.000 | 3836.5 | 9549.5 |
| BA.2.86.1 | USA | 0.788 | 0.778 | 0.797 | 1.001 | 3540.3 | 9338.5 |
| JN.1.31 | USA | 0.785 | 0.777 | 0.794 | 1.001 | 3024.5 | 7858.8 |
| JN.1.43 | USA | 0.783 | 0.776 | 0.789 | 1.002 | 1626.0 | 4370.9 |
| JN.1.45 | USA | 0.779 | 0.770 | 0.789 | 1.000 | 3632.5 | 9424.8 |
| JN.1.47 | USA | 0.773 | 0.763 | 0.782 | 1.000 | 4479.7 | 9930.4 |
| XBB.1.41.1 | USA | 0.770 | 0.754 | 0.784 | 1.000 | 10263.4 | 10552.7 |
| JN.1.24 | USA | 0.769 | 0.746 | 0.790 | 1.000 | 15822.4 | 11960.9 |
| JN.1.1 | USA | 0.764 | 0.760 | 0.769 | 1.003 | 741.7 | 2794.2 |
| JN.1.41 | USA | 0.752 | 0.738 | 0.765 | 1.000 | 9386.4 | 11016.5 |
| JN.3 | USA | 0.743 | 0.726 | 0.759 | 1.000 | 13184.5 | 11105.6 |
| GK.1.1 | USA | 0.742 | 0.719 | 0.764 | 1.000 | 17197.5 | 12248.1 |
| JN.2 | USA | 0.741 | 0.720 | 0.761 | 1.000 | 19978.5 | 11891.7 |
| EG.5.1.6 | USA | 0.733 | 0.708 | 0.756 | 1.001 | 19184.7 | 11226.6 |
| EG.5.1 | USA | 0.730 | 0.709 | 0.750 | 1.000 | 16758.3 | 12111.4 |
| JG.3 | USA | 0.729 | 0.720 | 0.737 | 1.000 | 3553.3 | 8529.4 |
| XBB.1.16.6 | USA | 0.715 | 0.694 | 0.735 | 1.000 | 17728.7 | 11911.2 |
| HK.3.2 | USA | 0.714 | 0.691 | 0.736 | 1.000 | 22929.1 | 11580.5 |
| EG.5.1.1 | USA | 0.713 | 0.692 | 0.732 | 1.000 | 21136.7 | 12282.2 |
| FL.1.5.1 | USA | 0.706 | 0.684 | 0.726 | 1.000 | 24029.5 | 11081.7 |
| JD.1.1.1 | USA | 0.706 | 0.685 | 0.725 | 1.000 | 19620.3 | 11755.3 |
| JD.1.1 | USA | 0.705 | 0.692 | 0.717 | 1.000 | 9400.7 | 10110.9 |
| HV.1 | USA | 0.704 | 0.698 | 0.710 | 1.001 | 1725.6 | 6366.0 |
| HK.3 | USA | 0.702 | 0.688 | 0.716 | 1.000 | 10173.5 | 10985.1 |
| XBB.1.16.17 | USA | 0.700 | 0.669 | 0.729 | 1.000 | 25174.4 | 12932.6 |
| FL.1.5.2 | USA | 0.698 | 0.660 | 0.732 | 1.000 | 21054.4 | 12042.6 |
| XBB.1.16.11 | USA | 0.696 | 0.663 | 0.727 | 1.000 | 24554.3 | 11098.6 |
| JF.1 | USA | 0.696 | 0.672 | 0.718 | 1.000 | 22331.0 | 11498.5 |
| XBB.1.16.15 | USA | 0.688 | 0.648 | 0.725 | 1.000 | 26191.9 | 12276.4 |
| XEC | United Kingdom | 1.108 | 1.088 | 1.129 | 1.000 | 23704.5 | 11595.1 |

| | | | | | | | |
|---|---|---|---|---|---|---|---|
| MC.1 | United Kingdom | 1.026 | 1.008 | 1.045 | 1.000 | 24382.6 | 11376.0 |
| MK.1 | United Kingdom | 0.962 | 0.944 | 0.981 | 1.000 | 19431.5 | 11535.7 |
| LW.1 | United Kingdom | 0.955 | 0.942 | 0.969 | 1.000 | 14816.4 | 11438.7 |
| KP.3.2.3 | United Kingdom | 0.943 | 0.930 | 0.957 | 1.000 | 12546.9 | 10265.1 |
| KP.2.14 | United Kingdom | 0.938 | 0.919 | 0.956 | 1.000 | 16156.8 | 10581.8 |
| KP.3.1.3 | United Kingdom | 0.935 | 0.923 | 0.948 | 1.000 | 9958.1 | 10021.3 |
| KP.1.1.3 | United Kingdom | 0.935 | 0.924 | 0.947 | 1.000 | 10093.9 | 10458.1 |
| KP.3.3 | United Kingdom | 0.929 | 0.923 | 0.935 | 1.001 | 3265.9 | 6972.1 |
| KP.2.6 | United Kingdom | 0.928 | 0.909 | 0.946 | 1.001 | 15937.4 | 10155.8 |
| LB.1.3 | United Kingdom | 0.921 | 0.908 | 0.934 | 1.001 | 8697.2 | 10436.5 |
| KP.3.2.2 | United Kingdom | 0.920 | 0.906 | 0.933 | 1.000 | 10231.1 | 10030.0 |
| KP.3.1 | United Kingdom | 0.919 | 0.914 | 0.924 | 1.002 | 1761.0 | 4680.4 |
| KP.3.2.4 | United Kingdom | 0.910 | 0.901 | 0.918 | 1.001 | 4343.9 | 8206.6 |
| KP.3.1.4 | United Kingdom | 0.907 | 0.898 | 0.916 | 1.001 | 4533.5 | 7387.4 |
| KP.2.3 | United Kingdom | 0.899 | 0.892 | 0.907 | 1.001 | 2980.7 | 6873.9 |
| JN.1.50 | United Kingdom | 0.895 | 0.878 | 0.912 | 1.000 | 11584.1 | 11112.1 |
| KP.2.2 | United Kingdom | 0.891 | 0.882 | 0.900 | 1.001 | 4128.2 | 7295.5 |
| JN.1.26 | United Kingdom | 0.886 | 0.868 | 0.904 | 1.000 | 12434.3 | 11717.5 |
| KP.2.8 | United Kingdom | 0.882 | 0.866 | 0.898 | 1.001 | 9276.6 | 10349.0 |
| KP.3.2 | United Kingdom | 0.877 | 0.870 | 0.884 | 1.001 | 2217.0 | 5350.2 |
| KP.3 | United Kingdom | 0.875 | 0.869 | 0.881 | 1.003 | 1575.8 | 3725.6 |
| JN.1.18.3 | United Kingdom | 0.869 | 0.851 | 0.887 | 1.000 | 10341.1 | 10579.4 |
| LB.1 | United Kingdom | 0.869 | 0.860 | 0.878 | 1.001 | 3483.2 | 6521.6 |
| KP.1.1 | United Kingdom | 0.852 | 0.844 | 0.859 | 1.002 | 2143.0 | 5311.4 |
| KP.1.1.1 | United Kingdom | 0.850 | 0.841 | 0.859 | 1.001 | 3077.5 | 5647.6 |
| LF.1 | United Kingdom | 0.849 | 0.834 | 0.865 | 1.001 | 7305.5 | 10031.5 |
| KP.2 | United Kingdom | 0.848 | 0.842 | 0.853 | 1.004 | 1117.7 | 3123.4 |
| KS.1 | United Kingdom | 0.847 | 0.839 | 0.855 | 1.001 | 2264.0 | 4970.2 |
| JN.1.10 | United Kingdom | 0.847 | 0.832 | 0.862 | 1.000 | 7077.9 | 8711.4 |
| JN.1.7.4 | United Kingdom | 0.847 | 0.835 | 0.858 | 1.001 | 4479.3 | 7715.3 |
| KP.4.1 | United Kingdom | 0.846 | 0.833 | 0.858 | 1.001 | 4951.4 | 8653.9 |
| JN.1.16.1 | United Kingdom | 0.835 | 0.829 | 0.842 | 1.003 | 1335.8 | 3825.4 |
| KW.1.1 | United Kingdom | 0.834 | 0.819 | 0.848 | 1.001 | 6551.9 | 8607.7 |
| LA.1 | United Kingdom | 0.834 | 0.822 | 0.845 | 1.001 | 4365.0 | 7641.1 |
| JN.1.13.1 | United Kingdom | 0.820 | 0.807 | 0.834 | 1.001 | 5132.2 | 8816.4 |
| XDK.1 | United Kingdom | 0.819 | 0.809 | 0.828 | 1.001 | 3184.6 | 5946.4 |
| JN.1.20 | United Kingdom | 0.819 | 0.807 | 0.830 | 1.001 | 4309.9 | 7071.9 |
| JN.1.8 | United Kingdom | 0.818 | 0.810 | 0.826 | 1.002 | 1927.2 | 4782.6 |
| JN.1.16 | United Kingdom | 0.815 | 0.808 | 0.822 | 1.003 | 1516.7 | 3733.8 |
| JN.1.32.1 | United Kingdom | 0.814 | 0.803 | 0.825 | 1.001 | 3736.6 | 7008.9 |

| | | | | | | | |
|---|---|---|---|---|---|---|---|
| JN.1.34 | United Kingdom | 0.799 | 0.788 | 0.811 | 1.001 | 3307.2 | 7445.4 |
| JN.1.7.1 | United Kingdom | 0.798 | 0.790 | 0.807 | 1.002 | 2009.7 | 4980.1 |
| JN.1.11.1 | United Kingdom | 0.797 | 0.788 | 0.806 | 1.002 | 2623.0 | 6489.0 |
| JN.1.7 | United Kingdom | 0.790 | 0.785 | 0.796 | 1.004 | 960.5 | 2322.8 |
| JN.1.4.4 | United Kingdom | 0.788 | 0.779 | 0.798 | 1.002 | 2482.1 | 6213.1 |
| JN.1.32 | United Kingdom | 0.784 | 0.777 | 0.791 | 1.004 | 1351.0 | 3638.2 |
| JN.1.7.2 | United Kingdom | 0.784 | 0.773 | 0.795 | 1.001 | 3478.7 | 7433.1 |
| JN.1.4.6 | United Kingdom | 0.782 | 0.774 | 0.791 | 1.003 | 2045.4 | 5870.8 |
| JN.1.18 | United Kingdom | 0.775 | 0.768 | 0.783 | 1.004 | 1452.3 | 4231.1 |
| XDK | United Kingdom | 0.773 | 0.765 | 0.782 | 1.002 | 1979.5 | 5510.0 |
| JN.1.9.1 | United Kingdom | 0.771 | 0.762 | 0.780 | 1.002 | 2139.9 | 5418.5 |
| JN.1.8.1 | United Kingdom | 0.767 | 0.756 | 0.777 | 1.001 | 3285.4 | 7042.7 |
| JN.1.6.1 | United Kingdom | 0.759 | 0.746 | 0.773 | 1.001 | 4553.4 | 9052.9 |
| JN.1.43 | United Kingdom | 0.758 | 0.746 | 0.770 | 1.001 | 4118.5 | 8231.3 |
| JN.1.5 | United Kingdom | 0.756 | 0.743 | 0.769 | 1.001 | 4675.2 | 8380.4 |
| JN.1.39 | United Kingdom | 0.752 | 0.744 | 0.759 | 1.003 | 1815.6 | 5104.9 |
| JN.1.9 | United Kingdom | 0.751 | 0.743 | 0.759 | 1.003 | 1890.6 | 4649.4 |
| JN.1.1.1 | United Kingdom | 0.741 | 0.726 | 0.756 | 1.001 | 6243.5 | 8671.2 |
| JN.1.4.5 | United Kingdom | 0.740 | 0.730 | 0.749 | 1.002 | 2629.6 | 6405.3 |
| JN.1.4 | United Kingdom | 0.737 | 0.731 | 0.744 | 1.004 | 1145.1 | 2752.7 |
| JN.1.65 | United Kingdom | 0.735 | 0.722 | 0.746 | 1.001 | 4458.6 | 8520.4 |
| JN.1.19 | United Kingdom | 0.733 | 0.721 | 0.745 | 1.001 | 3771.1 | 7105.5 |
| JN.1 | United Kingdom | 0.729 | 0.724 | 0.735 | 1.005 | 959.2 | 2470.9 |
| JN.1.4.7 | United Kingdom | 0.727 | 0.709 | 0.743 | 1.001 | 8418.9 | 10510.0 |
| JN.1.22 | United Kingdom | 0.723 | 0.712 | 0.734 | 1.001 | 3698.4 | 7920.7 |
| BA.2.86.1 | United Kingdom | 0.709 | 0.694 | 0.724 | 1.000 | 6527.2 | 9209.9 |
| JN.1.8.3 | United Kingdom | 0.709 | 0.686 | 0.729 | 1.001 | 10600.8 | 10145.0 |
| JN.1.2 | United Kingdom | 0.708 | 0.685 | 0.729 | 1.000 | 13166.7 | 11037.9 |
| JG.3 | United Kingdom | 0.702 | 0.681 | 0.722 | 1.000 | 10285.0 | 10727.8 |
| JN.6 | United Kingdom | 0.701 | 0.682 | 0.719 | 1.000 | 9308.8 | 10504.0 |
| XDN | United Kingdom | 0.698 | 0.683 | 0.713 | 1.001 | 6623.4 | 8896.8 |
| JN.1.1 | United Kingdom | 0.696 | 0.686 | 0.705 | 1.001 | 3068.0 | 6663.2 |
| JN.2 | United Kingdom | 0.693 | 0.677 | 0.709 | 1.001 | 8075.8 | 8352.9 |
| JN.3 | United Kingdom | 0.683 | 0.659 | 0.706 | 1.000 | 14986.8 | 11169.1 |
| HV.1 | United Kingdom | 0.660 | 0.628 | 0.690 | 1.001 | 17688.9 | 10598.7 |
| JD.1.1 | United Kingdom | 0.640 | 0.605 | 0.673 | 1.000 | 20325.9 | 11744.8 |
| XEC | France | 1.177 | 1.144 | 1.213 | 1.000 | 17094.1 | 11701.3 |
| MC.1 | France | 1.042 | 1.018 | 1.069 | 1.000 | 15181.1 | 11620.0 |
| XDY | France | 1.016 | 0.989 | 1.045 | 1.000 | 15130.6 | 11405.4 |
| KP.1.1.3 | France | 0.951 | 0.937 | 0.965 | 1.000 | 12263.1 | 11212.2 |

| | | | | | | | |
|---|---|---|---|---|---|---|---|
| KS.1.1 | France | 0.951 | 0.939 | 0.962 | 1.000 | 11112.6 | 10840.6 |
| JN.1.8 | France | 0.941 | 0.927 | 0.955 | 1.000 | 11226.2 | 11554.0 |
| KP.3.3 | France | 0.938 | 0.927 | 0.949 | 1.000 | 9756.5 | 10576.2 |
| KP.3.2.4 | France | 0.931 | 0.913 | 0.949 | 1.000 | 13602.1 | 11624.5 |
| KP.3.1 | France | 0.926 | 0.918 | 0.935 | 1.000 | 5858.1 | 8787.8 |
| KP.2.3 | France | 0.920 | 0.912 | 0.929 | 1.000 | 6224.6 | 9341.8 |
| KP.3 | France | 0.918 | 0.909 | 0.928 | 1.000 | 6310.4 | 10035.4 |
| LB.1.3 | France | 0.918 | 0.903 | 0.933 | 1.000 | 11527.3 | 11573.2 |
| KP.3.2 | France | 0.914 | 0.899 | 0.928 | 1.000 | 10214.2 | 11096.8 |
| KP.2.2 | France | 0.910 | 0.902 | 0.918 | 1.000 | 4899.0 | 8523.6 |
| KP.3.1.4 | France | 0.909 | 0.890 | 0.929 | 1.000 | 14031.5 | 12175.1 |
| JN.1.50.1 | France | 0.908 | 0.891 | 0.926 | 1.000 | 11613.9 | 11810.2 |
| JN.1.50 | France | 0.906 | 0.894 | 0.918 | 1.000 | 8433.8 | 10653.1 |
| LB.1 | France | 0.905 | 0.894 | 0.918 | 1.000 | 8507.1 | 10389.9 |
| KP.1.1 | France | 0.874 | 0.861 | 0.887 | 1.000 | 6075.6 | 8760.7 |
| KP.2 | France | 0.874 | 0.865 | 0.883 | 1.001 | 4111.8 | 8157.7 |
| KS.1 | France | 0.869 | 0.859 | 0.879 | 1.000 | 4268.7 | 7768.5 |
| KP.1.1.1 | France | 0.865 | 0.853 | 0.877 | 1.000 | 4708.3 | 8511.5 |
| XDK.4.1 | France | 0.862 | 0.849 | 0.876 | 1.001 | 5616.5 | 9371.7 |
| JN.1.16.1 | France | 0.857 | 0.848 | 0.866 | 1.001 | 2997.3 | 6440.9 |
| XDK.3 | France | 0.846 | 0.833 | 0.860 | 1.000 | 5258.9 | 8327.3 |
| JN.1.16 | France | 0.841 | 0.829 | 0.853 | 1.001 | 4083.8 | 7558.2 |
| JN.1.32 | France | 0.800 | 0.788 | 0.811 | 1.001 | 3134.0 | 6823.4 |
| JN.1.7 | France | 0.793 | 0.783 | 0.803 | 1.001 | 2502.8 | 5703.7 |
| JN.1.39 | France | 0.787 | 0.775 | 0.798 | 1.001 | 3012.8 | 5950.3 |
| XDK | France | 0.778 | 0.768 | 0.788 | 1.001 | 2355.5 | 5145.4 |
| JN.1.4 | France | 0.762 | 0.751 | 0.774 | 1.001 | 2855.4 | 6235.4 |
| JN.1 | France | 0.748 | 0.738 | 0.758 | 1.001 | 2341.2 | 5309.7 |
| JN.1.1 | France | 0.713 | 0.700 | 0.725 | 1.001 | 3696.8 | 8286.6 |
| XEC | Canada | 1.094 | 1.067 | 1.123 | 1.000 | 17354.8 | 10825.2 |
| MC.1 | Canada | 1.058 | 1.037 | 1.081 | 1.000 | 17380.9 | 11977.4 |
| XDY | Canada | 1.016 | 0.992 | 1.043 | 1.000 | 17332.5 | 11424.5 |
| MB.1.1 | Canada | 1.001 | 0.977 | 1.028 | 1.000 | 19027.7 | 10983.5 |
| KP.3.3.2 | Canada | 0.996 | 0.976 | 1.019 | 1.000 | 17320.7 | 11760.8 |
| KP.3.3.1 | Canada | 0.972 | 0.958 | 0.986 | 1.000 | 14626.9 | 9982.2 |
| KP.3.5 | Canada | 0.966 | 0.949 | 0.983 | 1.001 | 13211.1 | 11864.2 |
| LB.1.3 | Canada | 0.961 | 0.954 | 0.969 | 1.001 | 6197.5 | 11457.7 |
| KP.1.1.5 | Canada | 0.956 | 0.941 | 0.971 | 1.000 | 15252.6 | 12257.1 |
| LB.1.7 | Canada | 0.955 | 0.944 | 0.966 | 1.000 | 9439.8 | 11327.6 |
| LB.1.4 | Canada | 0.954 | 0.943 | 0.965 | 1.000 | 10218.6 | 11808.6 |

| | | | | | | | |
|---|---|---|---|---|---|---|---|
| KP.3.3.3 | Canada | 0.954 | 0.936 | 0.972 | 1.000 | 13332.2 | 12535.9 |
| MK.1 | Canada | 0.952 | 0.932 | 0.972 | 1.000 | 15474.4 | 12503.8 |
| JN.1.16.3 | Canada | 0.944 | 0.926 | 0.962 | 1.000 | 15860.9 | 11749.0 |
| LB.1.2 | Canada | 0.943 | 0.931 | 0.956 | 1.000 | 12514.3 | 11236.1 |
| JN.1.10 | Canada | 0.943 | 0.928 | 0.959 | 1.001 | 12990.0 | 11450.5 |
| LP.1 | Canada | 0.942 | 0.932 | 0.953 | 1.000 | 8790.3 | 10818.0 |
| JN.1.40 | Canada | 0.941 | 0.925 | 0.957 | 1.000 | 16552.3 | 11679.9 |
| KP.3.2.4 | Canada | 0.937 | 0.922 | 0.952 | 1.000 | 13725.0 | 11331.8 |
| KP.3.1.6 | Canada | 0.935 | 0.920 | 0.951 | 1.000 | 11908.3 | 11783.7 |
| KP.1.1.3 | Canada | 0.934 | 0.927 | 0.941 | 1.002 | 3619.1 | 8329.1 |
| KP.2.14 | Canada | 0.933 | 0.917 | 0.949 | 1.000 | 13621.9 | 11664.1 |
| KP.2.15 | Canada | 0.929 | 0.916 | 0.943 | 1.000 | 14139.4 | 10992.4 |
| JN.1.50 | Canada | 0.928 | 0.914 | 0.943 | 1.001 | 10703.4 | 10167.2 |
| KP.2.9 | Canada | 0.928 | 0.912 | 0.945 | 1.000 | 13059.5 | 11750.2 |
| KP.2.3 | Canada | 0.928 | 0.923 | 0.932 | 1.004 | 1585.4 | 4316.9 |
| KP.3.3 | Canada | 0.925 | 0.920 | 0.930 | 1.003 | 2225.8 | 5783.8 |
| KP.3.2.3 | Canada | 0.923 | 0.918 | 0.928 | 1.002 | 2128.4 | 5975.1 |
| KP.3.1.4 | Canada | 0.921 | 0.915 | 0.927 | 1.002 | 2872.3 | 7741.3 |
| KP.2.2 | Canada | 0.915 | 0.909 | 0.921 | 1.003 | 2789.2 | 6114.9 |
| LB.1.1 | Canada | 0.914 | 0.903 | 0.927 | 1.001 | 8792.6 | 10961.9 |
| KS.1.1 | Canada | 0.905 | 0.896 | 0.914 | 1.001 | 4891.1 | 8801.9 |
| KP.3.1 | Canada | 0.898 | 0.894 | 0.902 | 1.006 | 1132.6 | 2771.3 |
| LB.1 | Canada | 0.892 | 0.888 | 0.896 | 1.005 | 1156.6 | 3161.8 |
| XDV.1 | Canada | 0.890 | 0.880 | 0.899 | 1.001 | 5297.8 | 8880.3 |
| LF.2 | Canada | 0.881 | 0.870 | 0.893 | 1.001 | 8832.8 | 10697.8 |
| JN.1.7.4 | Canada | 0.879 | 0.865 | 0.892 | 1.000 | 8861.0 | 12379.0 |
| KP.3.2 | Canada | 0.879 | 0.874 | 0.883 | 1.005 | 1133.6 | 2755.2 |
| KP.1.1.1 | Canada | 0.875 | 0.869 | 0.880 | 1.003 | 2032.7 | 5384.9 |
| KP.3 | Canada | 0.871 | 0.867 | 0.875 | 1.007 | 958.3 | 2122.0 |
| JN.1.16.1 | Canada | 0.869 | 0.863 | 0.876 | 1.003 | 2356.3 | 6209.6 |
| KP.4.2 | Canada | 0.866 | 0.851 | 0.880 | 1.000 | 9829.8 | 11786.0 |
| KP.2 | Canada | 0.857 | 0.853 | 0.862 | 1.006 | 1081.5 | 2584.3 |
| KR.1 | Canada | 0.857 | 0.847 | 0.867 | 1.001 | 4584.0 | 9351.6 |
| KP.4.1 | Canada | 0.856 | 0.846 | 0.866 | 1.002 | 4134.8 | 6872.5 |
| JN.1.8 | Canada | 0.853 | 0.846 | 0.860 | 1.002 | 2414.7 | 5749.0 |
| KS.1 | Canada | 0.849 | 0.843 | 0.854 | 1.003 | 1743.8 | 4394.0 |
| KP.1.1 | Canada | 0.844 | 0.838 | 0.850 | 1.003 | 1903.1 | 4833.5 |
| JN.1.26 | Canada | 0.840 | 0.830 | 0.851 | 1.002 | 4820.9 | 9137.5 |
| KP.2.1 | Canada | 0.836 | 0.825 | 0.846 | 1.001 | 4882.5 | 9559.5 |
| JN.1.16 | Canada | 0.835 | 0.829 | 0.841 | 1.003 | 1909.2 | 4697.3 |

| | | | | | | | |
|---|---|---|---|---|---|---|---|
| JN.1.11.1 | Canada | 0.823 | 0.817 | 0.830 | 1.003 | 1921.4 | 4117.0 |
| JN.1.3 | Canada | 0.821 | 0.810 | 0.833 | 1.001 | 4847.7 | 7929.1 |
| JN.1.1.6 | Canada | 0.818 | 0.806 | 0.830 | 1.001 | 6060.9 | 9472.4 |
| XDK.1 | Canada | 0.817 | 0.808 | 0.826 | 1.002 | 3053.0 | 7031.5 |
| JN.1.34 | Canada | 0.814 | 0.804 | 0.823 | 1.001 | 3689.0 | 8193.8 |
| JN.1.20 | Canada | 0.813 | 0.802 | 0.825 | 1.001 | 5211.5 | 7766.2 |
| KP.1 | Canada | 0.808 | 0.797 | 0.818 | 1.001 | 4452.7 | 7700.4 |
| XDP | Canada | 0.805 | 0.798 | 0.811 | 1.003 | 1773.8 | 4788.5 |
| JN.1.32 | Canada | 0.802 | 0.795 | 0.809 | 1.003 | 1803.9 | 4355.2 |
| JN.1.13.1 | Canada | 0.801 | 0.796 | 0.807 | 1.004 | 1464.1 | 3510.6 |
| JN.1.7 | Canada | 0.801 | 0.796 | 0.806 | 1.007 | 956.9 | 2095.2 |
| XDK | Canada | 0.798 | 0.788 | 0.808 | 1.001 | 3677.4 | 8399.2 |
| JN.1.4.6 | Canada | 0.797 | 0.790 | 0.804 | 1.003 | 1784.5 | 4059.5 |
| JN.1.18 | Canada | 0.797 | 0.789 | 0.804 | 1.002 | 2131.2 | 5570.3 |
| JN.1.7.2 | Canada | 0.793 | 0.785 | 0.801 | 1.002 | 2562.5 | 6081.7 |
| JN.1.4.4 | Canada | 0.792 | 0.783 | 0.801 | 1.002 | 3075.7 | 6854.2 |
| JN.1.8.1 | Canada | 0.775 | 0.770 | 0.780 | 1.005 | 1084.8 | 2543.9 |
| JN.1.19 | Canada | 0.770 | 0.756 | 0.784 | 1.000 | 6830.3 | 11378.9 |
| JN.1.4 | Canada | 0.763 | 0.759 | 0.767 | 1.007 | 788.9 | 1706.4 |
| JN.1.46 | Canada | 0.756 | 0.742 | 0.770 | 1.001 | 6353.9 | 8716.0 |
| JN.1.39 | Canada | 0.755 | 0.747 | 0.763 | 1.003 | 2172.9 | 4981.2 |
| JN.1.4.7 | Canada | 0.749 | 0.736 | 0.763 | 1.001 | 7133.1 | 9441.8 |
| JN.1.4.5 | Canada | 0.749 | 0.743 | 0.755 | 1.004 | 1365.4 | 3894.6 |
| JN.1.5 | Canada | 0.743 | 0.731 | 0.755 | 1.001 | 5323.2 | 8494.5 |
| JN.1.9 | Canada | 0.743 | 0.734 | 0.752 | 1.002 | 3292.1 | 6372.6 |
| JN.1 | Canada | 0.743 | 0.739 | 0.747 | 1.008 | 744.9 | 1659.4 |
| JN.1.52 | Canada | 0.741 | 0.733 | 0.749 | 1.002 | 2343.9 | 5004.9 |
| JN.1.60 | Canada | 0.739 | 0.728 | 0.750 | 1.001 | 4328.7 | 8585.2 |
| JN.1.31 | Canada | 0.737 | 0.731 | 0.744 | 1.003 | 1809.1 | 4238.3 |
| JN.1.42 | Canada | 0.729 | 0.715 | 0.743 | 1.001 | 7249.8 | 9526.1 |
| JN.1.2 | Canada | 0.729 | 0.722 | 0.735 | 1.004 | 1722.2 | 4271.1 |
| JN.1.6 | Canada | 0.727 | 0.710 | 0.743 | 1.001 | 8560.6 | 11136.5 |
| JN.1.22 | Canada | 0.723 | 0.717 | 0.729 | 1.004 | 1480.5 | 4004.9 |
| JN.2.5 | Canada | 0.719 | 0.712 | 0.726 | 1.003 | 2166.3 | 4779.6 |
| JN.1.47 | Canada | 0.719 | 0.706 | 0.731 | 1.001 | 5438.2 | 8942.4 |
| JN.1.1 | Canada | 0.711 | 0.705 | 0.718 | 1.004 | 1705.7 | 4875.0 |
| JN.1.45 | Canada | 0.706 | 0.687 | 0.724 | 1.001 | 10726.5 | 10774.2 |
| EG.5.1.1 | Canada | 0.705 | 0.685 | 0.724 | 1.000 | 12219.6 | 11753.8 |
| JG.3 | Canada | 0.698 | 0.690 | 0.706 | 1.003 | 2636.0 | 6450.7 |
| BA.2.86.1 | Canada | 0.697 | 0.675 | 0.719 | 1.000 | 14402.4 | 11593.2 |

| | | | | | | | |
|---|---|---|---|---|---|---|---|
| JN.1.47.1 | Canada | 0.683 | 0.659 | 0.705 | 1.000 | 14000.1 | 11458.9 |
| JN.2 | Canada | 0.653 | 0.626 | 0.678 | 1.000 | 17291.0 | 11406.5 |
| HV.1 | Canada | 0.639 | 0.629 | 0.649 | 1.001 | 4680.3 | 8412.8 |
| JD.1.1 | Canada | 0.630 | 0.599 | 0.659 | 1.001 | 22077.0 | 11380.6 |
| HK.3 | Canada | 0.627 | 0.603 | 0.651 | 1.000 | 18133.5 | 10230.4 |
| JD.1.1.1 | Canada | 0.542 | 0.493 | 0.587 | 1.000 | 27210.7 | 11872.0 |
| XEC | Germany | 1.044 | 1.022 | 1.067 | 1.001 | 8913.2 | 9940.5 |
| KP.2.3 | Germany | 0.943 | 0.927 | 0.959 | 1.000 | 5213.1 | 7900.7 |
| KP.1.1.3 | Germany | 0.935 | 0.918 | 0.953 | 1.001 | 5497.8 | 7248.5 |
| KP.3.3 | Germany | 0.934 | 0.922 | 0.945 | 1.001 | 3182.5 | 5613.4 |
| KP.3.1 | Germany | 0.929 | 0.915 | 0.943 | 1.001 | 3935.5 | 6659.6 |
| KP.2 | Germany | 0.906 | 0.893 | 0.920 | 1.001 | 3160.9 | 6235.8 |
| JN.1.16 | Germany | 0.893 | 0.881 | 0.905 | 1.002 | 2847.1 | 5108.5 |
| JN.1.7 | Germany | 0.881 | 0.868 | 0.893 | 1.002 | 3045.7 | 5707.3 |
| JN.1 | Germany | 0.876 | 0.866 | 0.887 | 1.003 | 2326.7 | 4061.5 |
| JN.1.4 | Germany | 0.851 | 0.839 | 0.863 | 1.001 | 2997.9 | 5211.1 |
| JN.1.1 | Germany | 0.834 | 0.810 | 0.856 | 1.001 | 6861.4 | 8902.5 |

The relative Re of KP.3.1.1 is set to 1.

### Acknowledgments

We would like to thank all members of The Genotype to Phenotype Japan (G2P-Japan) Consortium. We thank Kenzo Tokunaga (National Institute of Infectious Diseases, Japan) for sharing materials, Jin Kuramochi (Interpark Kuramochi Clinic, Japan), Yuka Kamoshita (Department of Laboratory Medicine, Keio University School of Medicine, Japan), Masayo Noguchi (Clinical Laboratory, Keio University Hospital, Japan), Akatsuki Saito (Department of Veterinary Science, University of Miyazaki, Japan) and Chihiro Motozono, Yoshihiko Goto, Yuka Tajima, Takeshi Nakama, Kyoko Yamada (Division of Infection and Immunity, Kumamoto University, Japan) for supporting patient sera collection, and Mika Chiba, Kyoko Yasuda, Keiko Iida, Tsuki Fukuda (Division of Systems Virology, University of Tokyo, Japan) for performing experimental assays. We gratefully acknowledge the numerous laboratories worldwide that have provided sequence data and metadata to GISAID. A full list of originating and submitting laboratories for the sequences used in our analysis can be found at https://www.gisaid.org using the EPI-SET-ID: EPI_SET_241011ok and EPI_SET_241009gk.
